# Adenylate kinase-4 modulates oxidative stress and stabilizes HIF-1α to drive lung cancer metastasis

**DOI:** 10.1101/196857

**Authors:** Yi-Hua Jan, Tsung-Ching Lai, Chih-Jen Yang, Yuan-Feng Lin, Ming-Shyan Huang, Michael Hsiao

## Abstract

Disrupting signaling axes that are essential for tumor metastasis may provide therapeutic opportunity to cure cancer. We previously identified adenylate kinase 4 (AK4) as a biomarker of metastasis in lung cancer. Here we analyze AK4-associated metabolic gene signature and reveal HIF-1α is transcriptionally activated and associated with poor prognosis in lung adenocarcinoma patients. Overexpression of AK4 shifts metabolism towards aerobic glycolysis and elevates intracellular reactive oxygen species (ROS), which stabilizes and exaggerates HIF1-α protein expression and concurrently drives epithelial-to-mesenchymal transition (EMT) in hypoxia. Furthermore, overexpression of AK4 reduces hypoxic necrosis in tumors and promotes liver metastasis *in vivo*. Connectivity map analysis of AK4 gene signature identifies Withaferin-A as a potential compound to inhibit AK4-HIF-1α signaling axis, which then shows promising anti-metastatic potency in an orthotopic xenograft model of lung cancer. Our findings offer an alternative strategy to impair lung cancer metastasis via targeting AK4-HIF-1α axis.

## Introduction

Non-small cell lung cancer (NSCLC) remains the leading cause of cancer-related death around the world, mainly due to high metastatic rate (1). Recently, metabolic reprogramming has been considered as an important feature that drives the malignant progression of tumor (2). In metastatic cancer cells, energy metabolism is altered due to the constant exposure to oxidative stress and chronically depleted oxygen and nutrients. To fulfill biosynthetic and redox requirements, cancer cells consume glucose and secrete lactate even when oxygen is available, a phenomenon known as aerobic glycolysis or “Warburg effect” (3). Hypoxia-inducible factor-1α (HIF-1α) is a key transcription factor in response to hypoxic stress. HIF-1α transcribes genes involved in the process of glycolysis, angiogenesis and cancer metastasis (4,5). During metabolic stress, AMP-activated protein kinase (AMPK) is activated by sensing a decrease in the ratio of ATP to AMP, leading to the inhibition of ATP-consuming metabolic pathways and activates the energy-producing pathways (6). In addition, adenylate kinases (AKs), which are the abundant nucleotide phosphotransferases, catalyzing the generation of 2 molecules of ADP by transferring phosphate group from 1 molecule of ATP or GTP to AMP. The main role of AKs is to balance cellular adenine nucleotide composition to maintain energy homeostasis (7). However, the link between energy homeostasis and cancer progression has not been clearly elucidated.

Adenylate kinase 4 (AK4) is localized in mitochondrial matrix (8) and has been shown to physically bind to mitochondrial ADP/ATP translocase (ANT) as a stress-responsive protein to maintain cell survival (9). Moreover, several genomic and proteomic studies have shown AK4 expression fluctuates under cellular stress conditions (10-13). Significantly increased AK4 protein levels have been detected during development, in cultured cells exposed to hypoxia, and in an animal model of amyotrophic lateral sclerosis (9,14-16). Moreover, Lanning et al. showed that silencing of AK4 elevates cellular ATP level up to 25% and concurrently increases ADP/ATP ratio, which activates AMPK signalling (17). Previously, we identified AK4 as a lung cancer progression marker by assessing the correlation between AK4 levels and clinicopathological features (18). However, how AK4-iduced metabolic changes may affect cancer progression remains unclear.

Here we aimed to investigate the impact of AK4 expression on metabolic genes through analyzing lung cancer microarray datasets and decipher the functional consequences on lung cancer metastasis. We identified HIF-1α activity was significantly activated in AK4 metabolic gene signature in lung adenocarcinoma patients. Overexpression of AK4 shifts metabolism towards aerobic glycolysis and increases the levels of intracellular reactive oxygen species (ROS), which subsequently stabilizes HIF-1α protein and promotes epithelial-to-mesenchymal transition (EMT) of lung cancer cells in a HIF-1α-dependent manner. These findings represent a novel vicious cycle between AK4 and HIF-1α in response to hypoxic stress during lung cancer progression and highlighting the therapeutic opportunity of targeting AK4-HIF-1α axis in NSCLC.

## Materials and Methods

### Specimens

Clinical non-small cell lung cancer (NSCLC) samples were collected with IRB approval (KMUHIRB-E(I)-20160099) from the Kaohsiung Medical University Hospital and fixed in formalin and embedded in paraffin before archiving. Archived specimens, with follow-up time up to 200 months, were used for immunohistochemical staining. The histologic diagnosis was made according to the World Health Organization (WHO) classification guideline of lung cancer. The pathological diagnosis of tumor size, local invasion, lymph node involvement, distal metastasis and final disease stage were determined according to the American Joint Committee on Cancer (AJCC) TNM classification of lung cancer.

### Tissue microarray and immunohistochemical staining

Representative cores (1-mm-diameter) from each tumor sample were selected by matching histology from original hematoxylin and eosin (H&E)-stained slides and the histopathologic diagnosis of all samples were reviewed and confirmed by pathologists. IHC staining was performed using an automated immunostainer (Ventana Discovery XT autostainer, Ventana, USA) with 30-minute antigen retrieval procedure of heat-induced TRIS-EDTA buffer. Protein expressions were developed using 3, 3’-diaminobenzidine (DAB) peroxidase substrate kit (Ventana, USA). The following antibodies were used to detect AK4, HIF-1α, and E-cadherin protein expressions: AK4 (Genetex, 1:200), HIF-1α (Cell signaling, 1:100), and E-cadherin (Cell signaling, 1:100).

### Histology and IHC staining interpretation

The IHC staining results were assessed and scored independently by two pathologists who were blinded to patient’s clinical outcomes. Consensus decision was made when there was an inter-observer discrepancy. For scoring, both intensity and percentage of protein expression were recorded. The staining intensity was scored as: 0, no staining; 1+, weak staining; 2+, moderate staining; 3+, strong staining. The extent of staining was further divided into 2 groups by 25% of staining tumor cells. High IHC expression level was defined if staining intensity was 2+ or 3+ over 25% tumor cells.

### Microarray data analysis

The raw intensities from AK4 overexpression CL1-0 cells (GSE37930) and lung adenocarcinoma patient datasets (GSE31210) were normalized by robust multi-chip analysis (RMA) using GeneSpring GX11 (Agilent Technologies). AK4-associated gene signatures were identified by calculating the Pearson correlation coefficient between AK4 expression and each coding gene and ranked according to their correlation coefficient to AK4 expression. After applying Pearson correlation coefficient of ±0.3 as a threshold, AK4 metabolic gene signature was identified by selecting genes with enzyme or transporter annotations. Next, gene set enrichment analysis (GSEA) was performed to rank the probes and analyze gene set enrichment using c2.all.v5.1.symbols.gmt [curated] or c2.cp.kegg.v5.1.symbols.gmt [curated] gene sets as backend database (http://www.broadinstitute.org/gsea/). P-value less than 0.05 and FDR less than 25% were considered significant enrichment.

The activation or inhibition status of upstream regulators in AK4 metabolic gene signature was predicted using IPA Upstream Regulator Analysis (Ingenuity Systems, www.ingenuity.com) and the calculated z-scores can reflect the overall activation state of the regulator (<0: inhibited, >0: activated). In practice, Z-score more than 2 or less than -2 can be considered significant activation or inhibition, respectively.

### Cell lines

Human lung adenocarcinoma cell lines, H1355, PC9, H358, H928, CL1-0 CL1-1, CL1-3, and CL1-5, and squamous cell carcinoma, H157 and H520 were maintained in RPMI 1640 (Invitrogen) supplemented with 10% fetal bovine serum (FBS). Human lung adenocarcinoma cell lines (A549, PC13, and PC14) and large cell carcinoma, H1299 were grown in DMEM (Invitrogen) containing 10% FBS. All cells were kept under humidified atmosphere containing 5% CO2 at 37°C. CL1-0, CL1-1, CL1-3, and CL1-5 were established by Chu et al at National Taiwan University Hospital and displayed progressively increased invasiveness while PC13 and PC14 were derived from Tokyo National Cancer Centre Hospital. Other lung cancer cell lines (A549, H1355, H358, H928, H520, H157, H460, and H1299) were obtained from the American Type Culture Collection.

### Lentiviral shRNA and expression vectors

GIPZ Lentiviral AK4 (AK3L1) shRNA and HIF1A shRNA constructs, which carry the puromycin resistance gene, were purchased from Open Biosystems. Lentiviruses were generated by transfecting the shRNA-expressing vector and pMD2.G and pDeltaR8.9 into 293T cells using the calcium phosphate precipitation method. Virus-containing supernatants were collected, titrated, and used to infect cells with 8 μg/mL of polybrene. Infected cells were selected using 2 μg/mL of puromycin. For expression of AK4, full-length AK4 cDNA was cloned into pLenti6.3 lentiviral vector (Invitrogen). AK4-expressing cell lines were established by infecting cells with the pLenti6.3-AK4 viruses generated by transfecting pLenti6.3 AK4, pMD2.G and pDeltaR8.91 into 293T cells. Cells were then selected in 5 μg/mL of blasticidin.

### Western blot analysis

The following antibodies were used in western blot analysis: anti-AK4 (Genetex, 1:2,000), anti-HIF-1α (Cell signaling, 1:1,000), anti-hydroxylated HIF-1α (Cell signaling, 1:1,000), anti-E-cadherin (BD Bioscience, 1:1000), anti-vimentin (Sigma, 1:2,000), anti-Snail (Cell signaling, 1:1000) and anti–α-tubulin (Sigma-Aldrich, 1:5,000).

### Reagent and chemicals

Proscillaridin, Ouabain, Digitoxigenin, Digoxin, Withaferin-A, and Lanatoside-C were purchased from Sigma-Aldrich (St. Louis, MO). ATP Colorimetric Assay kit, glucose Colorimetric Assay kit, lactate Colorimetric Assay kit were purchased from BioVision (Milpitas, CA). CellROX Deep Red Reagent was purchased form Invitrogen.

### Cycloheximide assay

Cells were plated in 6-well plates and treated with cycloheximide (CHX) at a concentration of 50 μg/mL for 24 hours. Cells were then exposed to hypoxia for 6 hours to stabilize HIF-1α protein and then switched to normoxic condition. Protein lysates were harvested at 20-minute intervals in normoxic condition.

### ATP measurement

Cells were grown in 6-well plate overnight and the medium was refreshed with complete medium. After 24 hours, cells pellet was collected and the amount of ATP was quantified using ATP Colorimetric Assay kit (Biovision) according to manufacturer’s protocol.

### Glucose consumption assay

Cells were grown in 6-well plate overnight and the medium was refreshed with complete medium. After 24 hours, spent medium was collected and the amount of glucose was quantified using Glucose Colorimetric Assay kit (Biovision) according to manufacturer’s protocol.

### Lactate production assay

Cells were grown in 6-well plate overnight and the medium was refreshed with complete medium. After 24 hours, spent medium was collected and the amount of lactate was quantified using Lactate Colorimetric Assay kit (Biovision) according to manufacturer’s protocol.

### ROS measurement

ROS was quantified using CellROX Deep Red Reagent (Invitrogen).Briefly, cells were seeded in 96-well (2,000 cells/well) and were washed with PBS. Cells were then incubated with 5 μM of CellROX for 30 minutes at 37 °C and stained with DAPI. Intracellular ROS was measured using fluorescent plate reader with absorption/emission wavelength of ∼644/665 nm.

### Invasion assay

Polycarbonate filters were coated with human fibronectin on the lower side and Matrigel on the upper side. Medium containing 10% FCS was added to each well of the lower compartment of the chamber. Cells were suspended in serum-free medium containing 0.1% bovine serum albumin and loaded to each well of the upper chamber. After 16 hours, cells were fixed with methanol then stained Giemsa. Invaded cells were counted on lower side of the membrane under a light microscope (200×, ten random fields from each well). All experiments were performed in quadruplicate.

### Animal studies

All animal experiments were conducted according to protocols approved by the Academia Sinica Institutional Animal Care and Unitization Committee. Age-matched NOD-SCID Gamma (NSG) mice (6–8 weeks old) were used for xenograft model. For subcutaneous xenograft model, cells were subcutaneously injected into the flanks of NSG mice at the concentration of 1×10^6^ cells in 100 μL of PBS. Tumor volumes were measured weekly for 4 weeks. At the endpoint, tumors were removed and analyzed for hypoxic necrosis using H&E staining. Slides were digitally scanned using ScanScope AT (Aperio Technologies Inc.). Quantification of necrotic and non-necrotic area of H&E stained slides was performed using Definiens’ Tissue Studio software (Definiens Inc.). For the orthotopic xenograft model of lung cancer metastasis, 1×10^5^ cells (CL1-0 Vec or CL1-0 AK4) were suspended in 10 μl of PBS/matrigel mixture (1:1) and injected into left lung of NSG mice (n=6 per group). Mice from Withaferin-A-treatment group were administered i.p. with 1mg/kg body weight or 4mg/kg body weight of Withaferin-A in 100 μL of PBS three times per week. Control mice were injected with 100 μL of vehicle PBS consisting less than 10% of DMSO. Four weeks post-injection, mice were sacrificed and the metastatic liver nodules were counted by gross examination. H&E staining was performed to confirm the histology of metastatic nodules.

### Statistical analysis

Statistics analysis was performed on SPSS 17.0 software (SPSS, USA). The correlation of IHC expressions of AK4 and HIF-1α was assessed using Spearman’s rank correlation analysis. Estimates of the survival rates were analyzed by the Kaplan-Meier method and compared by log-rank test. For all analyses, a P-value <0.05 was considered statistically significant. All observations were confirmed by at least three independent experiments. The results are presented as the mean ± SD. We used two-tailed, unpaired Student’s t-tests for all pair-wise comparisons.

## Results

### HIF-1α transcription factor is active in AK4 metabolic gene signature in lung adenocarcinoma patients

To identify AK4 metabolic gene signature, we analyzed lung cancer dataset (GSE31210) that contains microarray data from 246 stage I/II lung adenocarcinoma patients. We first calculated the Pearson correlation coefficient between AK4 expression and each coding gene, and ranked according to their correlation coefficient to AK4 expression. We then determined AK4 metabolic gene signature by selecting genes annotated with enzymes or transporter and applying ±0.3 Pearson correlation coefficient as a cutoff threshold. By this mean, we identified 1,501 probes that are significantly associated with AK4 expression in lung adenocarcinoma patients (Fig. 1A). Next, we subjected AK4 metabolic gene signature to gene set enrichment analysis (GSEA). Interestingly, GSEA results showed that gene sets categorized in hypoxia response, HIF1A targets, glucose metabolism, and lung cancer prognostic genes were significantly enriched in AK4 metabolic gene signature (Fig. 1B). Furthermore, Ingenuity upstream regulator analysis showed HIF-1α was ranked as the top-one activated transcription factor with an activation Z-score of 4.3 (Figure 1C, left). The heatmap revealed both direct and indirect HIF-1α-regulated genes that were positively or negatively correlated with AK4 expression (Fig. 1C, right). Kaplan-Meier survival analysis showed that patients with high expression of AK4-HIF-1α signature were significantly associated with worse overall and relapse-free survival compared to patients with low level of AK4-HIF-1α signature (Fig. 1D). Meanwhile, we also identified the consensus AK4 metabolic gene signature between GSE31210 lung adenocarcinoma dataset and TCGA LUAD dataset and found HIF-1α was also significantly activated with an activation Z-score of 4.098 (Supplementary Fig. S1A and B).

**Figure 1.**
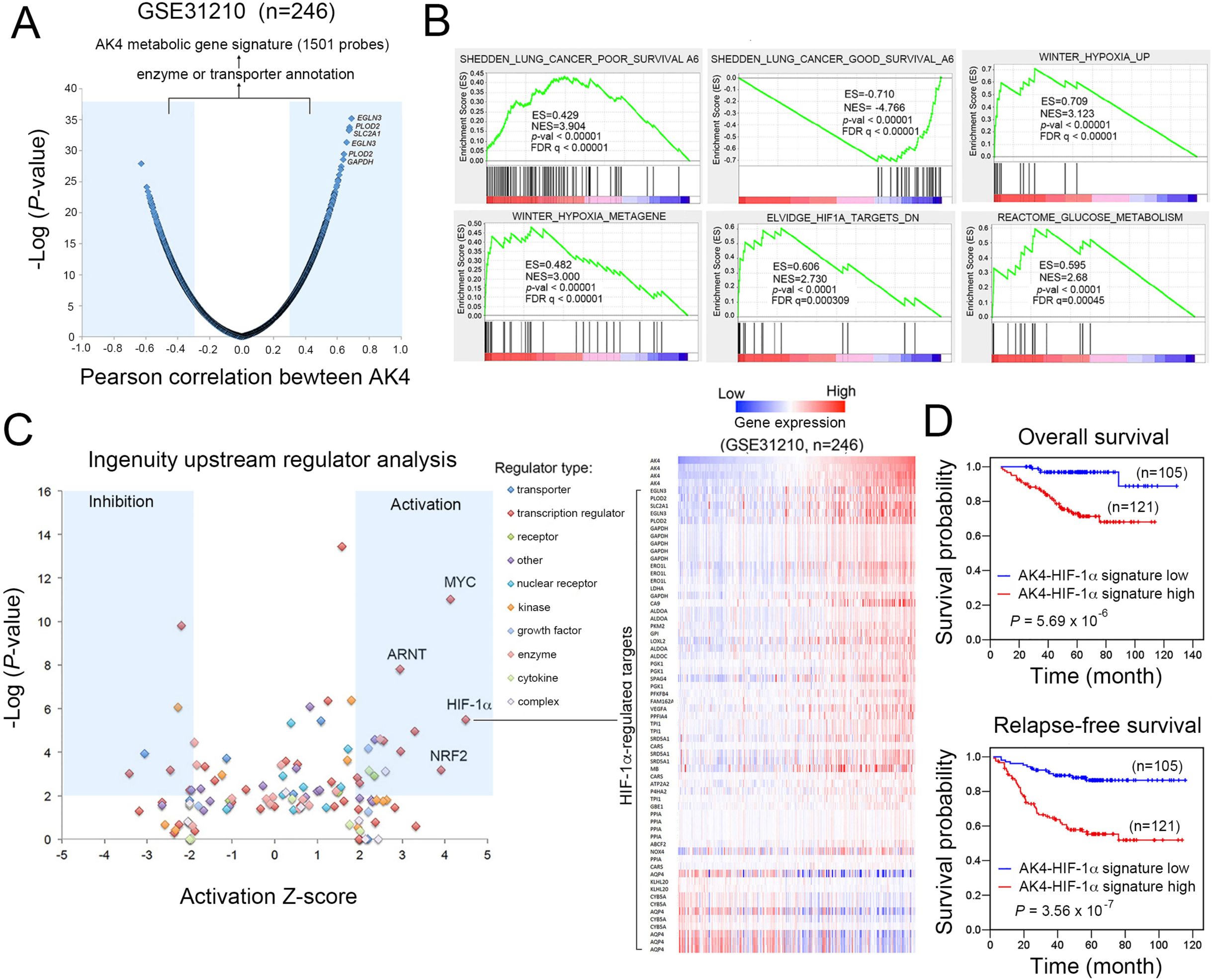
Upstream analysis of AK4 metabolic gene signature predicts HIF-1α is transcriptionally activated in lung adenocarcinomas. **A**, Genes were ranked according to their corresponding Pearson correlation coefficient (R) to AK4 expression. Genes that are positively correlated with AK4 (R ≥ 0.3) or negatively correlated with AK4 (R ≤ -0.3) are further filtered with enzyme or transporter annotations and defined as AK4 metabolic gene signature. **B**, GSEA plots of lung cancer prognostic genes, hypoxic response, and glucose metabolism in AK4 metabolic gene signature. **C**, Left, Ingenuity upstream regulator analysis algorithm predicts significant activation or inhibition of upstream regulators in AK4 metabolic gene signature. Activation Z-score more than 2 or less than -2 is considered to be significant activation or inhibition respectively. Right, heatmap illustrates both direct and indirect HIF-1α-regulated genes that positively or negatively correlate with AK4 expression. **D**, Left, overall survival analysis of patients stratified according to AK4-HIF-1α gene expression signature. Right, relapse-free survival analysis of patients stratified according to AK4-HIF-1α gene expression signature.

### Combination of AK4 and HIF-1α expression predicts worse prognosis compared to HIF-1 α alone in NSCLC patients

To validate the correlation between AK4 and HIF-1α in clinical specimens, we assessed their expression in 113 NSCLC patients by immunohistochemistry (IHC). Figure 2A showed the scoring criteria for quantifying the expression of AK4 and HIF-1α immunoreactivity in serial sections of IHC. Spearman correlation analysis of IHC results showed a significant positive correlation between AK4 and HIF-1α (Fig. 2B, Spearman’s rho= 0.457, *P* = 1,79 × 10^−6^). We next investigated the prognostic value of HIF-1α by IHC analysis and the results show patients with high HIF-1α tend to have worse survival compared to patients with low HIF-1α (Fig. 2C, left, *P*= 0.298). Furthermore, the combination of AK4 and HIF-1α status further revealed that patients with high AK4 and high HIF-1α exhibited significantly worse outcomes compared to the rest of the patients in this cohort (Fig. 2C, right, *P* = 0.017).

**Figure 2.**
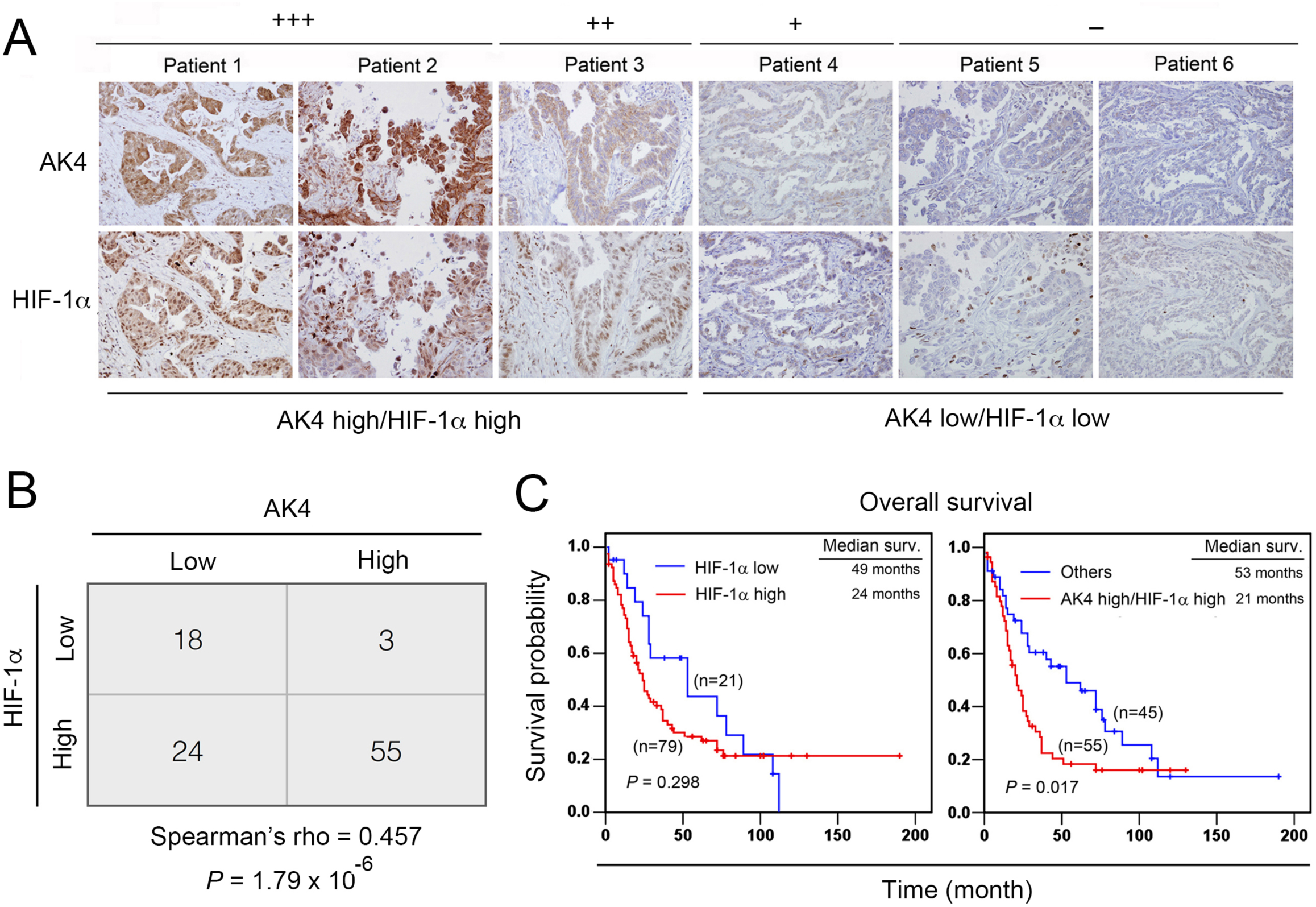
AK4 positively correlates with HIF-1α expression in NSCLC patients. **A**, Representative IHC images of AK4 and HIF-1α expression scores in serial sections of NSCLC tissues. **B**, Spearman’s rho correlation analysis of the IHC staining results for AK4 and HIF-1α in NSCLC patients. **C**, Overall survival analysis of 100 lung cancer patients stratified according to HIF-1α alone or combination of AK4 and HIF-1α.

### AK4 exaggerates HIF-1α protein in hypoxia and induces EMT

Next, we detected endogenous protein expression of AK4 and HIF-1α in 17 human NSCLC cell lines and found CL1-5, H441, H157, and CL1-3 cells expressed high levels of HIF-1α in normoxic condition (95% air, 5% CO2) and those cells also expressed with high levels of AK4 (Fig. 3A, upper). Correlation analysis showed a significant positive correlation between AK4 and HIF-1α (Fig. 3A, bottom). To assess the interaction between AK4 and HIF-1α, we knocked down endogenous AK4 expression using two AK4 shRNA (shAK4) clones in CL1-5. Interestingly, knockdown of AK4 suppressed HIF-1α with concurrent upregulation of E-cadherin and downregulation of vimentin (Fig. 3B, left). In a cell model of TGF-β-induced EMT, we found knockdown of AK4 partially hampered TGF-β-induced EMT with concurrent upregulation of phosphorylated AMPK at Thr172 (Fig. 3B, right). On the other hand, we overexpressed AK4 in CL1-0 cells and H1355 cells and exposed to hypoxic conditions (1% O2, 5% CO2, 94% N2) with different period of time. Western blot analysis showed that HIF-1α was upregulated earlier and to a higher degree in AK4-overexpressing cells compared to vector-expressing cells and concurrently induced EMT in CL1-0 and H1355 (Fig. 3C). We next investigated how AK4 regulates HIF-1α protein in CL1-0 cells. RT-PCR analysis showed that AK4 along with several HIF-1α downstream targets including VEGF, GLUT1, and HK2 were induced upon hypoxia. However, HIF-1α mRNA was not affected by AK4 overexpression (Fig. 3D). Therefore, we hypothesized the elevation of HIF-1α may be derived from translational control and/or post-translational regulation. We treated vector- and AK4-overexpressing CL1-0 cells with proteasome inhibitor MG-132 to block the protein degradation and detected HIF-1α protein under normoxia and hypoxia and found the effect of AK4 overexpression on HIF-1α protein elevation was diminished when proteasome was inhibited (Fig. 3E). To test whether the enhancement of HIF-1α by AK4 is through protein stabilization, we exposed the vector- and AK4-overexpressing CL1-0 cells to hypoxia to induce HIF-1α protein and then treated the cells with cycloheximide (CHX) to block de novo protein synthesis and assessed HIF-1α protein levels at 20 minutes interval under normoxic condition. We found that the HIF-1α protein was significantly stabilized in the AK4-overexpressing cells compared to the vector control cells (Fig. 3F). Moreover, we also found that the AK4-overexpressing cells had less hydroylated HIF-1α indicating the prolyl hydroxylase (PHD) activity is lower in the AK4-overexpressing cells (Fig. 3G). Moreover, overexpression of AK4 promoted invasion activity of CL1-0 and H1355 cells under hypoxia (Fig. 3H and I). To test whether HIF-1α played a critical role in AK4-induced EMT and invasion activity, we inhibited HIF-1α expression by shRNA in in AK4-overexpressing and vector-expressing CL1-0 cells. Knockdown of HIF1A in AK4-overexpressing cells partially suppressed EMT and suppressed AK4-induced invasion activity under hypoxia (Supplementary Fig S2A and B).

**Figure 3.**
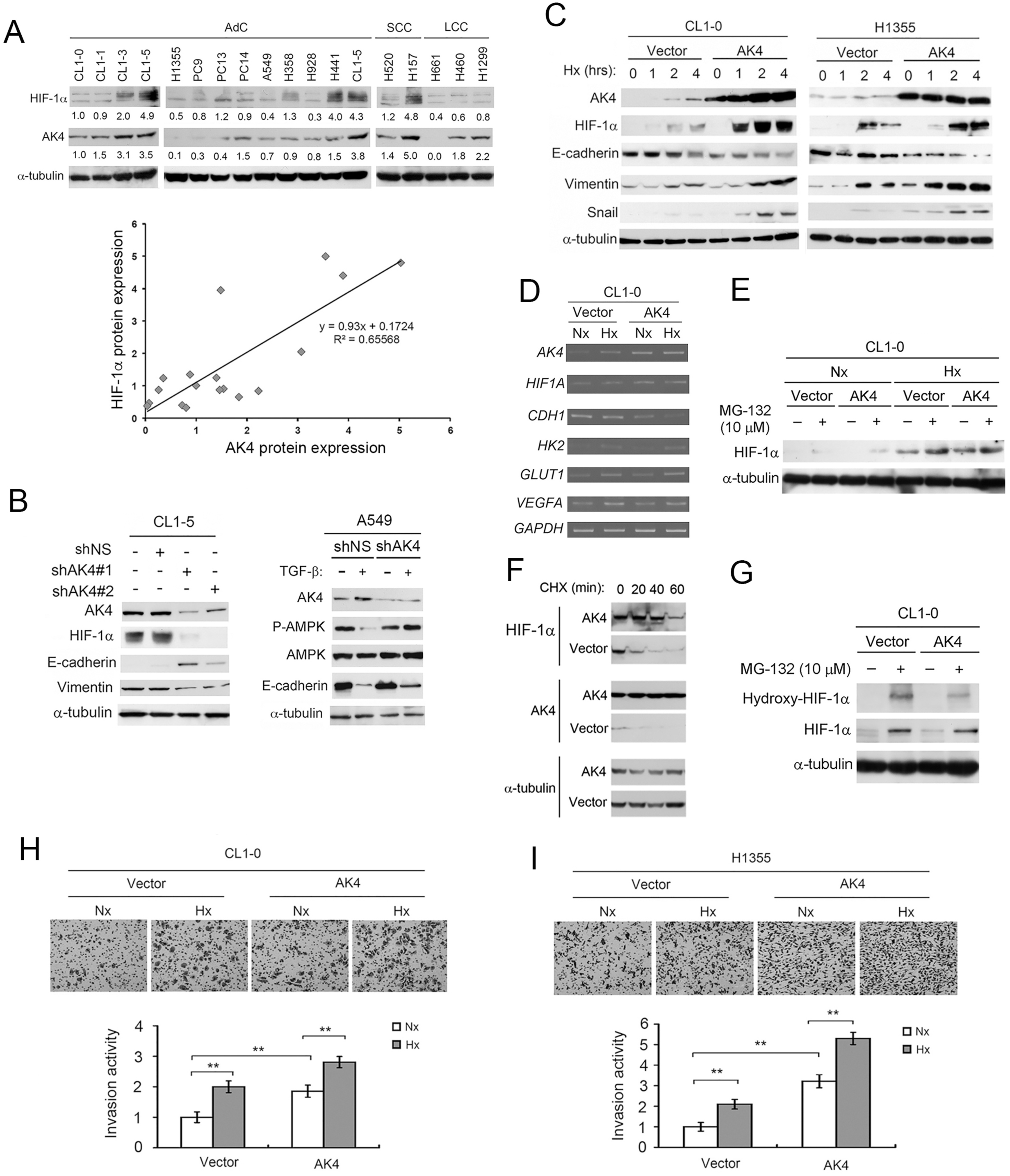
AK4 stabilizes and exaggerates HIF-1α protein expression to promote EMT. **A**, Upper, endogenous AK4 and HIF-1α protein expression in human NSCLC cell lines. Bottom, correlation between AK4 and HIF-1α protein expression in NSCLC cell lines. **B,** Left, WB analysis of AK4, HIF-1α, E-cadherin, Vimentin upon AK4 knockdown in CL1-5 cells. Right, WB analysis of AK4, AMPK, Phospho-AMPK (Thr172), and E-cadherin upon AK4 knockdown in A549 cells treated with or without TGF-β (5 ng/mL) for 24 hours. **C**, WB analysis of HIF-1α, AK4, E-cadherin, Vimentin, Snail from CL1-0 and H1355 vector- or AK4-overexpressing cells exposed to hypoxia for indicated time. **D**, RT-PCR analysis of *AK4*, *HIF1A*, *CDH1*, *HK2*, *GLUT1*, *VEGFA* and *GAPDH* in CL1-0 vector- or AK4-expressing cells under normoxic and hypoxic condition. **E**, WB analysis of HIF-1α from CL1-0 vector- or AK4-expressing cells treated with proteasome inhibitor MG-132 under Nx and Hx conditions. **F**, WB analysis of HIF-1α and AK4 from CL1-0 vector- or AK4-expressing cells treated with CHX for 20, 40, and 60 min. **G**, WB analysis of HIF-1α and hydroxylated HIF-1α from CL1-0 vector or AK4-expressing cells treated with MG-132 in Nx. **H**, Invasion assay of CL1-0 vector- or AK4-expressing cells in normoxia (Nx) or hypoxia (Hx). **P<0.01. **I**, Invasion assay of H1355 vector- or AK4-expressing cells in Nx or Hx. \*\**P*<0.01.

### AK4 elevates intracellular ROS and promotes aerobic glycolysis

To decipher the possible metabolic pathways affected by AK4, GSEA was used to analyze GSE37930 microarray data of differentially expressed genes (1.5 fold change) of CL1-0 AK4 cells versus CL1-0 Vec cells. We found 12 KEGG gene sets were positively enriched at the threshold of p-value<0.05 and FDR<25%. The enriched gene sets that positively correlate with AK4 can be further categorized into four super metabolic pathways including carbohydrate, amino acids, xenobiotics and oxidative stress (Fig. 4A). Notably, upregulation of AK4 was significantly correlated with genes in glycolysis_gluconeogenesis metabolism and glutathione metabolism (Fig. 4B; Supplementary Fig. S3A and B). These data prompted us to investigate the effect of AK4 overexpression on glycolysis and oxidative stress. By measuring levels of ATP, glucose and lactate, we found that overexpression of AK resulted in increased ATP and glucose consumption and increased lactate production (Fig. 4C). Moreover, overexpression of AK4 significantly decreased ADP/ATP ratio compared to control upon hypoxia (Fig. 4D). To quantify ROS levels within the cell, we used CellROX deep red to probe intracellular ROS levels in AK4-expressing and vector-expressing CL1-0 cells. The result showed overexpression of AK4 increased ROS levels to 1.67 fold compared to control (Fig. 4E). Furthermore, antioxidant N-acetyl-cysteine (NAC) treatment abolished AK4-induced stabilization of HIF-1α and invasion under hypoxia (Fig. 4F and G). On the other hands, knockdown of AK4 in CL1-5 cells reduced ROS levels nearly 20 percent compared to shNS control (Fig. 4H). NAC treatment in CL1-5 cells not only decreased HIF-1α and AK4 protein in a time-dependent fashion but also inhibited its invasion activity (Fig. 4I and J).

**Figure 4.**
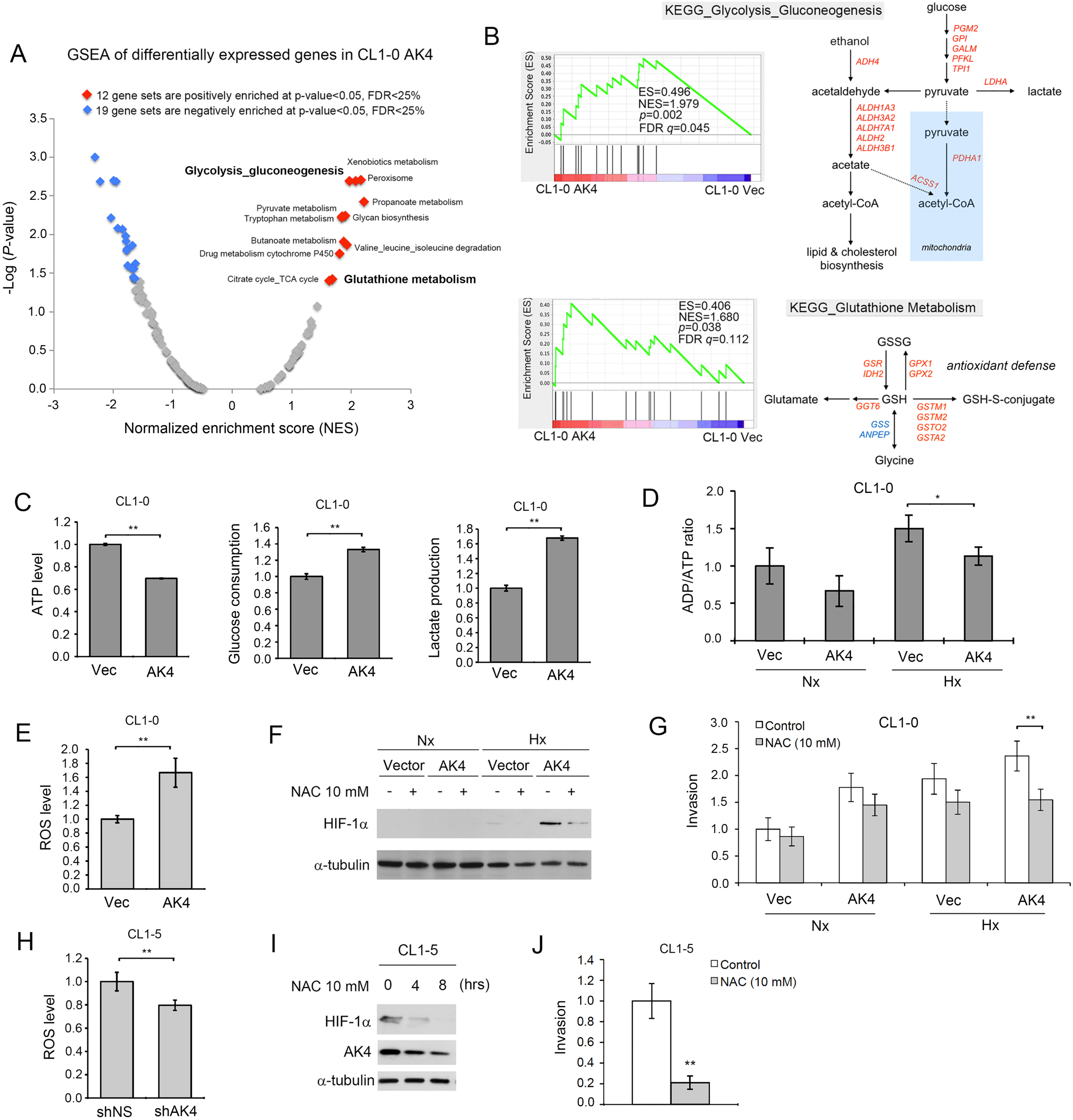
AK4 shifts metabolism towards aerobic glycolysis and increases oxidative stress. **A**, Global GSEA statistics of differentially expressed genes in AK4-overexpressing CL1-0 cells compared to vector expressing CL1-0 cells. **B**, GSEA plots and KEGG metabolic pathways of Glycolysis_Gluconeogenesis and Glutathione_Metabolism pathways between CL1-0 AK4 cells and CL1-0 Vec cells. **C**, Relative ATP levels, glucose consumption, and lactate production upon AK4 overexpression in CL1-0 cells. **P<0.01 **D**, Relative ADP/ATP ratio upon AK4 overexpression in CL1-0 cells under Nx or Hx. **E**, Intracellular ROS level of vector-expressing CL1-0 cells and AK4-expressing CL1-0 cells, ROS level was normalized to vector-expressing CL1-0 cells. \*\**P*<0.01. **F**, WB analysis of HIF-1α in CL1-0 vector- and AK4-expressing cells treated with or without 10 mM of NAC under Nx or Hx. **G**, Invasion assay of CL1-0 vector- or AK4-expressing cells treated with 10 mM of NAC under Nx or Hx. **H**, Intracellular ROS level of shNS-expressing CL1-5 cells and shAK4-expressing CL1-5 cells. ROS level was normalized to shNS-expressing CL1-5 cells. \*\**P*<0.01. **I**, Time course analysis of HIF-1α and AK4 protein expression in CL1-5 cells treated with 10 mM of NAC for indicated time. **J**, Invasion assay of CL1-5 cells treated with or without 10 mM of NAC.

### AK4 reduces hypoxic necrosis and promotes metastasis *in vivo*

We next investigated the effects of AK4 overexpression on tumorigenecity and metastasis *in vivo*. CL1-0 vector- and AK-expressing cells were subcutaneously injected in NSG mice. Four weeks post-injection, there was no significant differences in terms of tumor volume between CL1-0 Vec tumors and CL1-0 AK4 tumors (Fig. 5A). However, by examining H&E staining, we surprisingly found the necrotic tumor area was dramatically reduced in CL1-0 AK4 tumors compared to CL1-0 Vec tumors (Fig. 5B). We further performed IHC analysis of serial sections from CL1-0 AK4 and CL1-0 Vec xenograft tumors and revealed a strong positive correlation between AK4 expression and nuclear HIF-1α and a negative correlation between AK4 and E-cadherin in vivo (Fig. 5C). To further determine the effect of AK4 overexpression on metastasis, we injected AK4-expressing and vector-expressing CL1-0 cells into left lung of NSG mice. Four weeks post-injection, we found overexpression of AK4 significantly promoted CL1-0 cells to metastasize to liver (Fig. 5D).Quantification of the metastatic nodules using H&E staining and histological examination confirmed the number of nodules in liver was significantly increased in the mice carrying AK4-overexpressing tumors compared to vector-expressing tumor (Fig. 5E).

**Figure 5.**
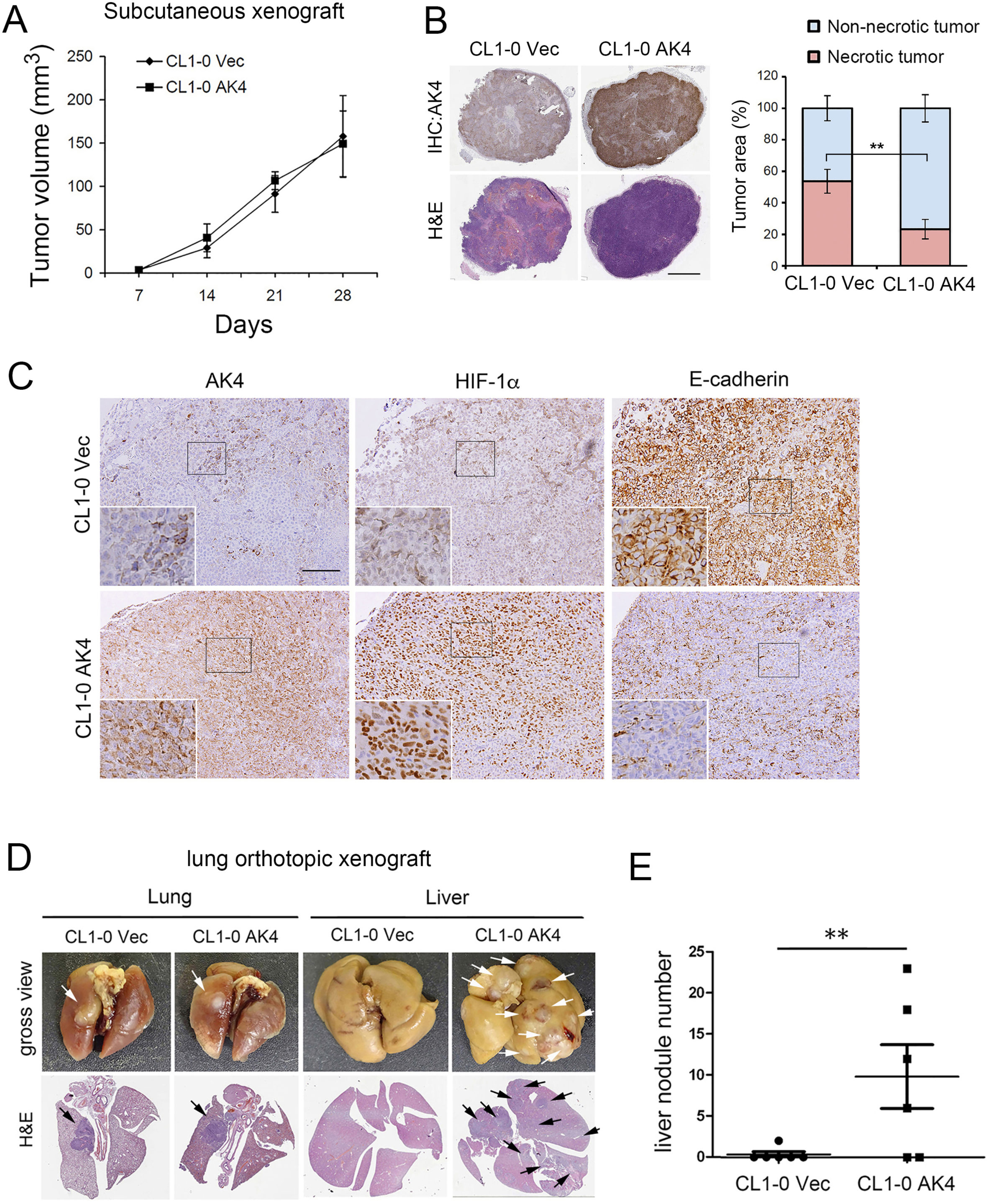
AK4 overexpression reduces hypoxic necrosis and promotes liver metastasis *in vivo*. **A**, Left, NOD Scid Gamma (NSG) mice were injected subcutaneously with CL1-0 Vec and CL1-0 AK4 cells (1x106 cells/100 μL) at left and right flanks respectively. Right, tumor volumes of CL1-0 Vec and CL1-0 AK4 were measured weekly as indicated. **B**, Left, H&E staining and IHC images of AK4 expression in CL1-0 Vec and CL1-0 AK4 subcutaneous xenograft tumors. Scale bar represents 2mm. Right, necrotic tumor area was detected and quantified by Definiens imaging analysis algorithm. \*\**P* <0.01. **C**, Representative IHC staining for AK4, HIF-1α, and E-cadherin expression in subcutaneous xenograft tumors from CL1-0 vector- or CL1-0 AK4-cells. Scale bar represents 100 μm. **D**, NSG mice were injected orthotopically at left lung with CL1-0 Vec or CL1-0 AK4 cells at the concentration of 1x10^5^ cells in 10 μL of PBS/matrigel mixture. **E**, Gross view (formalin-fixed) and H&E staining images in the lungs and livers of mice orthotopically injected with CL1-0 Vec or CL1-0 AK4 cells at day 30. The white and black arrows indicate tumor nodules in gross view and H&E staining images respectively. **F**, Quantification of liver nodule number of the mice orthotopically injected with CL1-0 Vec or CL1-0 AK4 cells at day 30. \*\**P*<0.01

### Identification of novel drug candidates for metastatic lung cancer through querying pharmacogenomics profiles of AK4 gene signature

To identify drug candidates that could inverse AK4 gene expression profiles as therapeutic strategy to inhibit lung cancer metastasis, we queried connectivity map using differentially expressed genes upon AK4 overexpression in CL1-0 and identified six structurally similar candidate drugs that showed the best enrichment scores including Proscillaridin, Ouabain, Digitoxigenin, Digoxin, Withaferin-A, and Lanatoside-C (Fig. 6A). We then conducted MTT cell viability assays after exposure to the drugs in culture and confirmed IC_50_ and IC_10_ for each drugs in CL1-0, CL1-5, CL1-0 Vec and CL1-0 AK4 (Supplementary Fig. S4). Furthermore, we screened the anti-invasion effect of each drug at the dose of IC_10_ in A549 and CL1-5 cells and found Withaferin-A, Lanatoside-C and Digoxin significantly suppressed invasion ability of A549 and CL1-5 cells (Fig. 6B). Interestingly, Digoxin and Lanatoside-C were proven to be potent HIF-1 modulators while the effect of Withaferin-A (WFA) on modulating HIF-1 α and AK4 was not clear. We then treated A549, CL1-5, CL1-0 Vec and CL1-0 AK4 cells with WFA in Nx or Hx and WB analysis showed HIF-1α and AK4 protein levels were significantly down-regulated upon WFA treatment (Fig. 6C). To evaluate the anti-metastatic effect of WFA, we orthotopically injected CL1-0 AK4 cells into left lung of NSG mice and the mice were subsequently treated with WFA at the dose of 1.0 mg/kg and 4.0 mg/kg three times per week through intraperitoneal administration. At day 28, the lungs and livers were removed for pathological examinations. The quantitative data of liver nodule number confirmed the anti-metastatic effect of WFA as mice received WFA at 1.0 mg/kg showed significant reduction of liver metastatic nodules (Fig. 6D and E). In mice received WFA at 4.0 mg/kg, the data showed promising therapeutic effect of suppressing both primary and metastatic tumors (Fig. 6D and E). Similar anti-metastatic effect of WFA was also observed in treating A549 orthotopic lung cancer model as mice received WFA at 1.0 mg/kg displayed significant inhibition of liver metastasis while mice received WFA at 4.0 mg/kg showed the lowest lung tumor burden with no sign of distal metastasis (Supplementary Fig. S5A-S5C). Taken together, our data suggest AK4-HIF-1α signaling axis is a potential therapeutic target of lung cancer metastasis and WFA might be a potential compound for further development to treat metastastic lung cancer (Fig. 6F).

**Figure 6.**
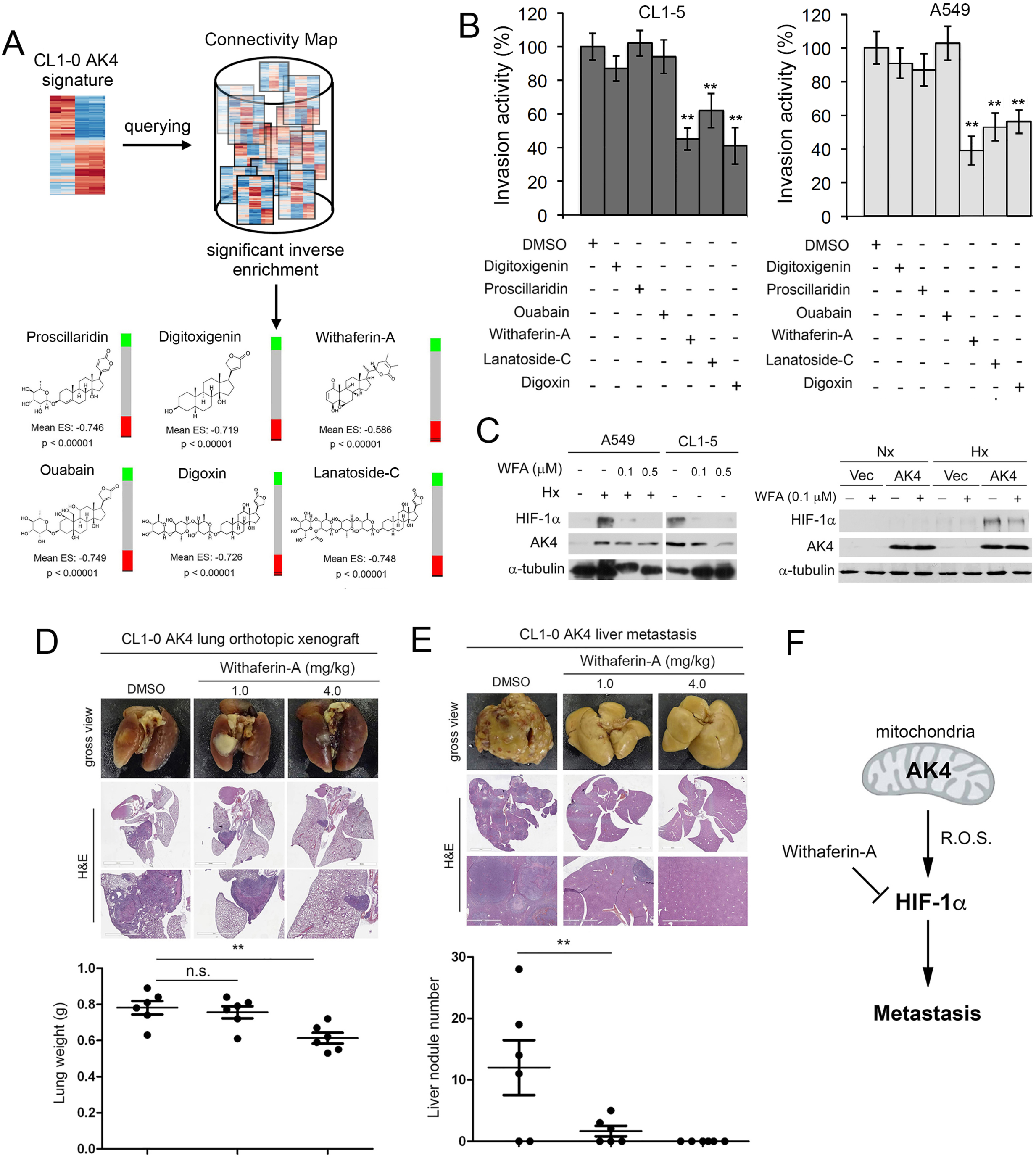
Connectivity map analysis of AK4 gene signature identifies Withaferin-A as a potent anti-metastatic agent for NSCLC. **A**, Identification of structurally similar drug candidates with best inverse AK4 gene expression enrichment score by querying connectivity map. **B**, Invasion assay of A549 and CL1-5 cells treated with corresponding IC10 doses of drug candidates. Data were expressed as percent inhibition compared to DMSO vehicle control. \*\**P* <0.01 **C**, WB analysis of HIF-1α and AK4 protein levels in A549, CL1-5, CL1-0 vector- and AK4-expressing cells treated with or without Withaferin-A under Nx or Hx. **D**, Gross view (formalin-fixed) and H&E staining images of the lungs in mice treated with DMSO vehicle control or Withaferin-A (1.0 mg/kg or 4.0 mg/kg) at day 30 after orthotopic injection of CL1-0 cells overexpressed with AK4 (*top*). Quantification of tumor weight in the lung of mice treated with DMSO vehicle control or Withaferin-A (1.0 mg/kg or 4.0 mg/kg) at day 28 after orthotopic injection of CL1-0 cells overexpressed with AK4 (*bottom*). Data are presented as the means ± SD. **, *P* < 0.01. **E**, Gross view (formalin-fixed) and H&E staining images of the livers in mice treated with DMSO vehicle control or Withaferin-A (1.0 mg/kg or 4.0 mg/kg) at day 30 after orthotopic injection of CL1-0 cells overexpressed with AK4 (top). Quantification of liver nodule number of the mice treated with DMSO vehicle control or Withaferin-A (1.0 mg/kg or 4.0 mg/kg) at day 30 after orthotopic injection of CL1-0 cells overexpressed with AK4 (bottom). Data are presented as the means ± SD. **, P < 0.01. **F**, Diagram depicts the working model of AK4-induced HIF-1α stabilization via elevating intracellular ROS, leading to subsequent EMT and metastasis. Targeting AK4-HIF-1α axis by withafein-A impairs lung cancer metastasis.

## Discussion

Dysregulation of HIF is increasingly recognized as a critical step during cancer progression (19). Deletion of *Hif1a* has been reported to markedly impair metastasis in mouse mammary tumor virus (MMTV) promoter-driven polyoma middle T antigen mouse model of breast cancer (20). In orthotopic xenograft model of lung cancer, HIF-1α antagonist PX-478 effectively inhibits tumor progression (21). Here we identified a novel signaling axis whereby enhanced expression of AK4 exaggerates HIF-1α protein expression, leading to EMT induction in lung cancer. Previous studies have shown that AK4 is one of the hypoxia responsive genes and AK4 is also a transcriptional target of HIF-1α (22-25). Surprisingly, we found the presence of AK4 can regulate exert feedback regulation of HIF-1α and this AK4-HIF-1α positive feedback loop is operative through the elevation of intracellular ROS which stabilizes HIF-1α protein and induces EMT.

In lung cancer, both small cell and non–small cell lung cancers express high levels of HIF-1α but its role as a prognostic factor remains controversial. HIF-1α expression has been reported as a poor prognosis marker in several studies (26-29). However, some studies reported inconsistent results showing that the predictive power of HIF-1α as prognosis marker is only marginal (30-32). In our study, patients with high levels of HIF-1α showed the trend towards poor prognosis compared to those with low levels of HIF-1α. Furthermore, we found combination of AK4 and HIF-1α status could significantly augment the prognostic power compared to HIF-1α alone (Fig. 2C). Taken together, our results suggest AK4 may serve as a critical factor for dictating the prognostic power of HIF-1α in lung cancer patients.

It is widely known that the regulation of HIF-1α by is mainly at the level of protein stability. In normoxic condition, HIF-1α is hydroxylated at two conserved proline residues (P402 and P577) by a family of HIF prolyl hydroxylase enzymes, including PHD1, PHD2 and PHD3. Hydroxylated HIF-1α is then polyubiquitinated by E3 ubiquitin ligase leading to proteosomal degradation. In hypoxia, hydroxylation does not occur due to the lack of substrate oxygen for PHDs. Moreover, ROS have been reported to inhibit the activity of PHDs (33-36). In our study, overexpression of AK4 reduced HIF-1α hydroxylation in the presence of MG132, suggesting that AK4 may stabilize HIF-1α protein by decreasing PHDs activity via ROS. Although the mechanism of AK4-mediated ROS production is unclear, the subcellular localization and physiological function of AK4 may provide a clue to explain the possible mechanism. AK4 interacts with ANT and voltage-dependent anion channel (VDAC) at mitochondrial matrix, and their interactions are required for the regulation of mitochondria membrane permeability and the export of ATP from mitochondrial matrix to cytosol in exchange for the import of ADP from cytosol to mitochondrial matrix (9). It is possible to hypothesize that AK4 may regulate the efflux of ROS generated from the electron transport chain (ETC) to cytosol through interacting with ANT/VDAC complex. Our results are consistent with other studies suggesting that ROS generated from the ETC could contribute to HIF-1α stabilization through blocking HIF-1α hydroxylation and von Hippel Lindau (pVHL) protein binding (37-39).

Prior studies have reported that hypoxia or overexpression of HIF-1α can induce EMT through direct binding of HIF-1α to hypoxia response element (HRE) within Snail and Twist promoter (40,41). Other EMT regulators such as Zeb1, Zeb2, and TCF-3 have been reported to be upregulated in pVHL-null renal cell carcinoma in which HIF-1α is constitutively overexpressed (42). In addition to binding to canonical HRE, HIF-1α can also interact with varieties of co-factors to activate EMT-associated genes and diverse gene expression in response to hypoxia (43,44).

Tumors rewire metabolism to provide sufficient energy and biosynthetic intermediates to meet the requirements for uncontrolled proliferation and progression. Enhanced glucose metabolism not only produces energy but also generates provision of macromolecular precursors and maintains NADPH homeostasis for cancer cells to withstand oxidative stress (45). Recent study suggested increased production of ROS is essential to enable and sustain metastatic phenotype (46). However, large-scale clinical trials using antioxidant supplement as preventive and therapeutic anticancer strategy did not show beneficial effect for cancer patients. On the contrary, antioxidant supplement even increased tumor incidence in genetically engineered mouse model of lung cancer and melanoma (47,48). One possible explanation for this controversial is that cancer cells adapt to have a tight redox regulation that allow them to withstand higher ROS accumulation than normal cells but below a critical cytotoxic threshold. The use of general antioxidants might alleviate circulating tumor cells from oxidative stress and accelerated metastasis development. Therefore, building up oxidative stress and equipped with antioxidant defenses is critical for tumor to metastasize (49). By ingenuity upstream regulator analysis of AK4 metabolic signature, we also identified NRF2, the master regulator of antioxidant responses, was significantly activated suggesting that lung adenocarcinoma patients with high AK4 expression may accompany with NRF2 activation (Fig. 1C). Furthermore, microarray analysis revealed genes encode for enzymes in glutathione metabolism pathway were differentially expressed upon AK4 overexpression in CL1-0 cells (Fig. 4B; Supplementary Fig. S3B). Moreover, in animal studies, we showed overexpression of AK4 not only protects tumor from hypoxic necrosis but also enhances its ability to metastasize. These data are consistent with the notion that only cancer cells equipped with enhanced antioxidant defense are capable of leveraging oxidative stress to promote metastasis. Our findings suggest overexpression of AK4 may trigger metabolic adaptation toward increased intracellular oxidative stress and antioxidant capacity at the same time and subsequently promote HIF-1α-mediated EMT and metastatic dissemination.

By pharmacogenomics analysis, we identified withaferin-A as a potential inhibitor that inverses AK4 induced gene signatures and it acts as a potent anti-metastatic agent in lung cancer. Similar to our findings, Hahm *et al*. showed withaferin-A treatment inhibits incidence of lung metastasis through suppressing glycolysis in mouse mammary tumor virus–neu (MMTV-neu) transgenic model, which highlights the therapeutic opportunities for targeting metabolic vulnerability of tumors (50).

In conclusion, we suggest that overexpression of AK4 stabilizes HIF-1α protein through increasing intracellular ROS and induces EMT in NSCLC. More importantly, pharmacologically inversing AK4 gene signature (e.g. by Withaferin-A) may serve as an effective strategy to treat metastatic lung cancer.

## Acknowledgement

This research was supported by Academia Sinica and Ministry of Science and Technology [MOST 104-0210-01-09-02,MOST 105-0210-01-13-01, MOST 106-0210-01-15-02, and MOST 105-2320-B-001-027-MY3]. We also thank Miss Tracy Tsai for her excellent works in immunohistochemistry staining analysis.

